# Representations of tactile object location in the retrosplenial cortex

**DOI:** 10.1101/2022.12.06.519323

**Authors:** Andreas Sigstad Lande, Koen Vervaeke

## Abstract

Little is known about how animals use tactile sensation to detect important objects and remember their location in a world-based coordinate system. Here, we hypothesized that retrosplenial cortex (RSC), a key network for contextual memory and spatial navigation, represents the location of objects based on tactile sensation. We studied mice that palpate objects with their whiskers while running on a treadmill in a tactile virtual reality in darkness. Using two-photon Ca^2+^ imaging, we discovered a population of neurons in agranular RSC that signal the location of tactile objects. Tactile object location responses do not simply reflect the sensory stimulus. Instead, they are highly task- and context-dependent and often predict the upcoming object before it is within reach. In addition, most tactile object location neurons also maintain a memory trace of the object’s location. These data show that RSC encodes the location and arrangement of tactile objects in a spatial reference frame.

## Introduction

Humans and animals effortlessly remember the location of important objects in their surroundings. To locate objects, humans mostly rely on their visual sense1. In addition, tactile interaction with objects contributes to remembering their location (Gallace and Spence, 2014), and for nocturnal animals such as rodents, tactile exploration is essential. However, how the brain encodes the location of objects based on tactile input remains unknown.

Functional imaging experiments of sighted and blind human subjects, who explore and memorize spatial layouts using only tactile input from the fingers (Wolbers et al., 2011; Ottink et al., 2021), show prominent activity in two brain areas: the parahippocampal- and the retrosplenial cortex (RSC)(Wolbers et al., 2011). The latter area may be surprising because RSC is best known for encoding visual information. Indeed, because of its prominent anatomical connections with the visual system and hippocampus (Sugar et al., 2011; Wyss and Groen, 1992; Groen and Wyss, 1992), RSC plays a crucial role in contextual memory and navigation when visual landmarks are available (Vann et al., 2009; Claessen and Ham, 2017; Maguire, 2001; Chen et al., 1994; Alexander and Nitz, 2015; Fischer et al., 2020; Mao et al., 2017; Vedder et al., 2016; Czajkowski et al., 2014; Cowansage et al., 2014; Bar, 2004; Powell et al., 2020; Mao et al., 2018; Rice et al., 1986). The experiments in humans suggest that RSC also has access to tactile information and may form a sensory modality-independent representation of space (Wolbers et al., 2011). However, testing this idea requires using a non-human animal model, such as rodents, to enable cellular-resolution circuit analysis during well-controlled behaviors.

To explore tactile objects, rodents use their facial whiskers - specialized hairs extending from follicles packed with nerve endings (Rice et al., 1986; O’Connor et al., 2021). Decades of research have revealed how the whisker system encodes the location of objects in the rodents’ peri-personal space in a head-centred coordinate system (Diamond et al., 2008; Petersen, 2007; Knutsen and Ahissar, 2009; Pluta et al., 2017; Pammer et al., 2013; O’Connor et al., 2010; Cheung et al., 2020). However, animals also need to remember the location of tactile objects in an extended space, in a world-based coordinate system. To date, no studies have examined how or where in the brain this representation is generated. Tactile information is likely routed to the brain’s spatial network and integrated with the spatial code (Gener et al., 2013; Save et al., 1998; O’Keefe and Dostrovsky, 1971). In support of this, whisker-related neural activity is not restricted to somatosensory areas but can also be observed in spatially modulated brain areas (Mohan et al., 2019; Olcese et al., 2013; Gallero-Salas et al., 2021; Mohan et al., 2018b, a; Bellistri et al., 2013; Pereira et al., 2007; Itskov et al., 2011; Merre et al., 2018). However, previous investigations were largely restricted to measuring neural responses to whisker stimuli in immobile or anaesthetized mice. Therefore, it remains to be tested how tactile information is processed during spatial navigation, and which circuits encode the location of tactile objects.

Here, we designed a task whereby mice run on a treadmill through a whisker-based tactile virtual reality system (inspired by (Sofroniew et al., 2014; Ayaz et al., 2019)). We found that neurons in agranular RSC represent the spatial location where the whiskers contact an object. These tactile object location responses show several features compatible with the hypothesis that RSC constructs a map of tactile space. Tactile object location responses did not simply reflect the sensory stimulus. Instead, they were highly task- and visual context-dependent. Moreover, in well-trained mice, a subset of tactile object location neurons also predicts of the upcoming object, and most object locations cells maintain their spatial tuning when the object is omitted. These data indicate that RSC encodes the location of tactile objects in a world-based spatial reference frame.

## Results

### Responses to tactile objects during a spatial task in darkness

We trained head-fixed mice to run laps on a treadmill in darkness to find the location of a water reward. At two positions along the track, a motorised cue, controlled in a closed loop by running speed, brushed across the whiskers, simulating the sensation of running past an object (Figure 1A, see Methods). Our goal was to develop a spatial behavioural task whereby we could change the tactile stimuli flexibly in a lap-by-lap manner and determine the precise timing of contacts between the whiskers and the object (Figure 1B. We performed these experiments in complete darkness to avoid visual responses to the moving object and therefore we tracked the animals’ behaviour using infrared light (940 nm) (Couto et al., 2019; Nikbakht and Diamond, 2021). Well-trained mice ran at a relatively constant speed, uninterrupted by the tactile stimulus, but slowed down to lick for water at the correct location, indicating they learned the rewarded position (Figure 1C, Video S1, data from 8 mice, 10 sessions).

**Figure 1.**
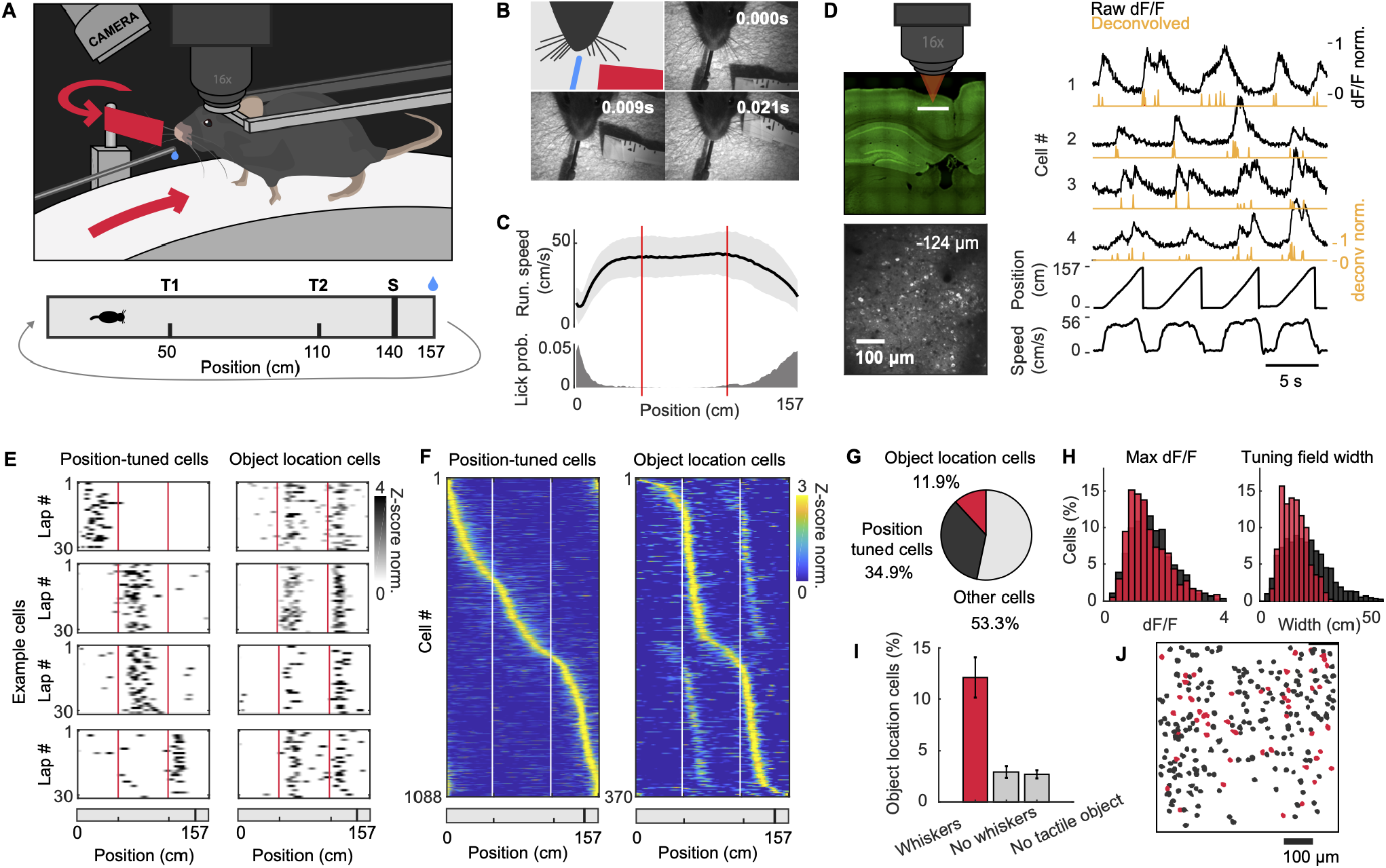
Responses to tactile objects during a spatial task in darkness. (**A**) Head-fixed mouse running laps in darkness to find the location of a water reward in a tactile virtual reality. T1 and T2 are the locations along the track where a tactile cue, controlled in a closed loop by running speed, brushes across the whiskers, simulating the sensation of running past an object. S is a 1 cm long sandpaper strip on the running wheel just before the reward location. (**B**) Whisker imaging using infrared (940 nm) illumination. The first frame is a cartoon outlining the mouse’s head and whiskers (black), waterspout (blue), and tactile object (red). (**C**) (Top) Mean ±SD running speed. (Bottom) Lick probability (8 mice, 10 sessions). (**D**) (Top) Confocal fluorescence image of coronal brain slice and microscope objective above agranular RSC (Thy1-GCaMP6s / GP 4.3 mice). (Bottom) Two-photon fluorescence image showing a typical field of view (FOV) of layer 2/3 neurons in agranular RSC (124 *μ*m deep). (Right) Example fluorescence traces (fractional change of fluorescence, dF/F) during four laps, and bottom traces showing the mouse’s position and running speed. Orange traces show deconvolved dF/F. (**E**) Examples of four position-tuned (left column) and four object location-tuned cells (right column). Deconvolved activity for 30 consecutive laps. Red lines indicate the movement onset of the tactile object. (**F**) Average response of all position-tuned cells and object location-tuned cells, sorted according to the position of peak response. White lines indicate the movement onset of the tactile object (8 mice, 10 FOVs). (**G**) Percentage of cell types (8 mice, 10 FOVs, 3120 cells in total, 1088 position-tuned cells, 370 object location cells). (**H**) (Left) Distribution of the maximum amplitude (dF/F) for all position-tuned cells (black) and object location cells (red). Right: Distribution of tuning curve width (8 mice, 10 FOVs, 1088 position-tuned cells, 370 object location cells). Kolmogorov-Smirnoff test for max dF/F: p = 4 × 10-4, and for tuning curve width: p = 2.7 × 10-19. (**I**) Percentage of object location cells (mean ± SEM) in: (1) Control mice exposed to the tactile object (“whiskers”, 8 mice, 10 FOVs, 370 out of 3120 cells), (2) in mice exposed to tactile objects but trained without contralateral whiskers (“no whiskers”, 3 mice, 6 FOVs, 40 cells out of 1448 cells), and (3) in mice with intact whiskers but without tactile objects (“no tactile object”, 2 mice, 3 FOVs, 20 out of 734 cells). (**J**) Spatial organization of cells in a typical FOV. Only position-tuned (black) and object location cells (red) are shown (184 position-tuned cells and 70 object location cells).

To measure neuronal activity in layer 2/3 of the agranular RSC (Brodmann area 30), contralateral to the stimulated whiskers, we performed two-photon Ca^2+^ imaging using mice expressing the Ca^2+^ indicator GCaMP6s in excitatory neurons (Thy1-GCaMP6s, 8 mice, 10 Fields of View, FOV) (Dana et al., 2014)(Figure 1D, see Methods). We deconvolved the fluorescence signals to infer spiking activity (Giovannucci et al., 2019). When we analyzed all neurons with stable firing fields in specific positions along the track, we observed two types of activity. Neurons with a single firing field that could be located anywhere along the track, which we refer to as position-tuned cells (Mao et al., 2017), and neurons with two firing fields tightly coupled to location of the tactile objects which we refer to as object location cells (34.9% position-tuned cells, 11.9% object location cells, out of 3120 neurons in total, Figure 1E-G). The response amplitude did not differ much between these groups, but the tuning width of object location cells was slightly narrower compared to position-tuned cells (Figure 1H). In mice trained without whiskers, the percentage of object location cells was greatly reduced and was similar to the percentage observed by chance in mice that had whiskers but were trained without tactile objects (Figure 1I, “Whiskers”: 12.1 ± 2 %, 8 mice, 10 FOVs; “No whiskers” 2.9 ± 0.6 %, 3 mice, 6 FOVs; “No tactile object”; 2.7 ± 0.4 %, 2 mice, 3 FOVs; mean ± SEM and hereafter, see Methods). We also considered whether whisker stimuli could evoke eye movements(Zahler et al., 2021) and whether these could trigger neuronal responses in RSC (Sikes et al., 1988)(Figure S1A-C, 3 mice, 3 FOVs). Tracking the pupils showed that tactile stimuli did not evoke any eye movements along the yaw or pitch rotation axis. However, we noticed that in some laps the mouse makes a minute eyeblink following a tactile stimulus (Figure S1D). We tested whether the activity of object location neurons was significantly correlated with these eyeblinks. However, this was the case for only 4.4 % of these cells showing that object location tuning does not depend on eye blinks (Figure S1E). Finally, when we visualized the spatial organization of object location neurons, we found that these were spatially intermingled with position-tuned neurons in a salt-and-pepper fashion suggesting that RSC may integrate object location and animal position signals (Figure 1J). In summary, these data demonstrate that a population of neurons encoding the location of tactile objects co-exist with position-tuned cells in agranular RSC.

### Tactile object location representations are task specific

Neurons in hippocampus and association cortices such as RSC encode not only the location of the animal but also what the animal is doing. Such a conjunctive code can disambiguate sensory stimuli that happened in the same environment but when the animal was engaged in different behaviors (Fischer et al., 2020; Pho et al., 2019; Franco and Goard, 2021; Kentros et al., 2004; Harvey et al., 2012; Miller et al., 2019). To test whether object location neurons in RSC are task-specific, we measured how the same neurons respond to tactile stimuli under three different task conditions. First, mice started the session performing the tactile virtual reality task (“goal oriented”) (Figure 2A). Next, we immobilized the mouse by clamping the running wheel and we provided playback tactile stimuli (“passive stimulus”, see Methods). Finally, we unclamped the running wheel to let mice run freely, but without water rewards, and we provided tactile objects in random locations along the track (“unengaged”).

**Figure 2.**
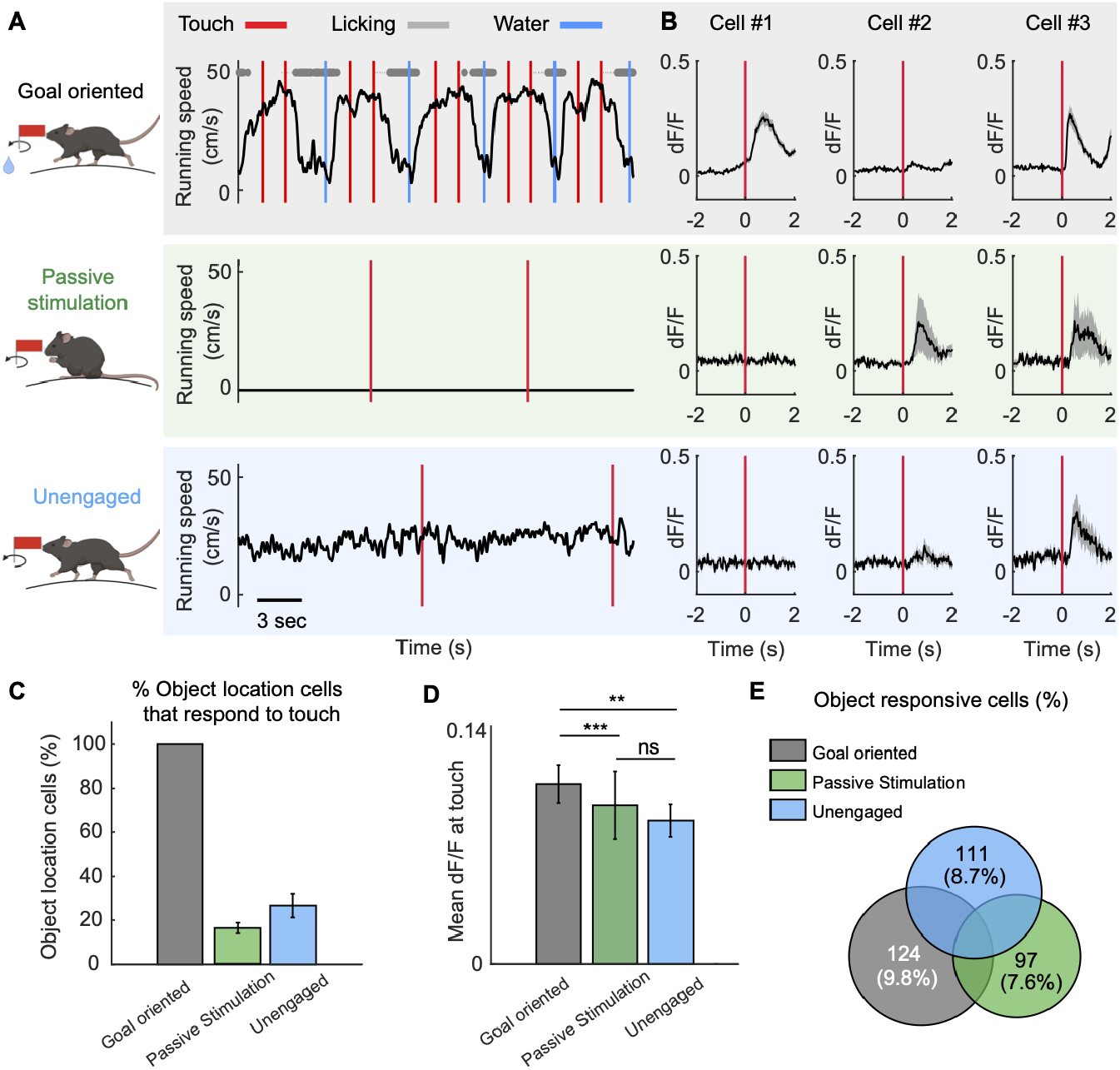
Responses to tactile objects are task specific. (**A**)(Left) Cartoons showing the three different task conditions. (Top) “Goal oriented”: Mice ran laps to find water rewards while encountering tactile objects along the path. (Middle) “Passive stimulation”: The running wheel was clamped to immobilize the mouse and playback tactile stimuli were provided at random time intervals separated by at least 5 seconds. (Bottom) “Unengaged”: Mice were free to run but there were no rewards, and tactile stimuli occurred at random time intervals. (**B**) Example cells illustrating the diversity of responses (mean ± SEM). Left example cell responded only in the goal-oriented task. The other two example cells responded under multiple task conditions. (**C**) Percentage of cells classified as object location cells underthegoal-oriented condition that were also responsive to tactile stimuli under the othertaskconditions (3 mice, 4 FOVs, 1271 cell recorded in total, “goal-oriented”, 100 %; “passive stimulation”, 16.1 ± 2.4 %; “unengaged”, 26.6 ± 5.4 %). (**D**) Touch response amplitude under each task condition (mean ± SEM, Mann Whitney-U test, “ns” p-value = 0.48; ** p-value = 0.02; *** p-value = 7.7 ·10^−4^). (**E**) Venn diagram showing the percentage of object-responsive cells under different task conditions.

How neurons responded to tactile objects was greatly task-dependent (Figure 2B). Of all object location neurons classified in the goal-oriented task (124 out of 1271 cells), only 16.1 2.4 % responded to passive stimuli, and 26.6 ± 5.4 % responded when the mouse was unengaged (Figure 2B, C, 3 mice, 4 FOVs). Overall, the response amplitude was only slightly different between conditions. Yet, in the goal-oriented task, object responses were significantly larger than responses under the other two task conditions (Figure 2D). Notably, the percentage of responsive neurons was similar under each task condition (Figure 2E). In total, 20 % of all neurons responded to tactile objects in at least one condition, showing that a large fraction of RSC neurons can respond to tactile objects but only do so under specific task conditions.

### Responses to tactile objects are visual context dependent

RSC may encode an abstract representation of a tactile object within its environment rather than of the tactile object per se (Bar, 2004; Zhao et al., 2020; Fyhn et al., 2007). Because of RSC’s well-established role in processing visual information, we hypothesized that tactile responses in RSC may be modulated by visual context. To test this, we presented tactile objects in different levels of ambient luminance such that the mouse can both see and feel the object. A session was composed of three blocks of laps (Figure 3A). The first block of laps occurred in darkness, followed by two blocks of laps in two different levels of ambient luminance.

**Figure 3.**
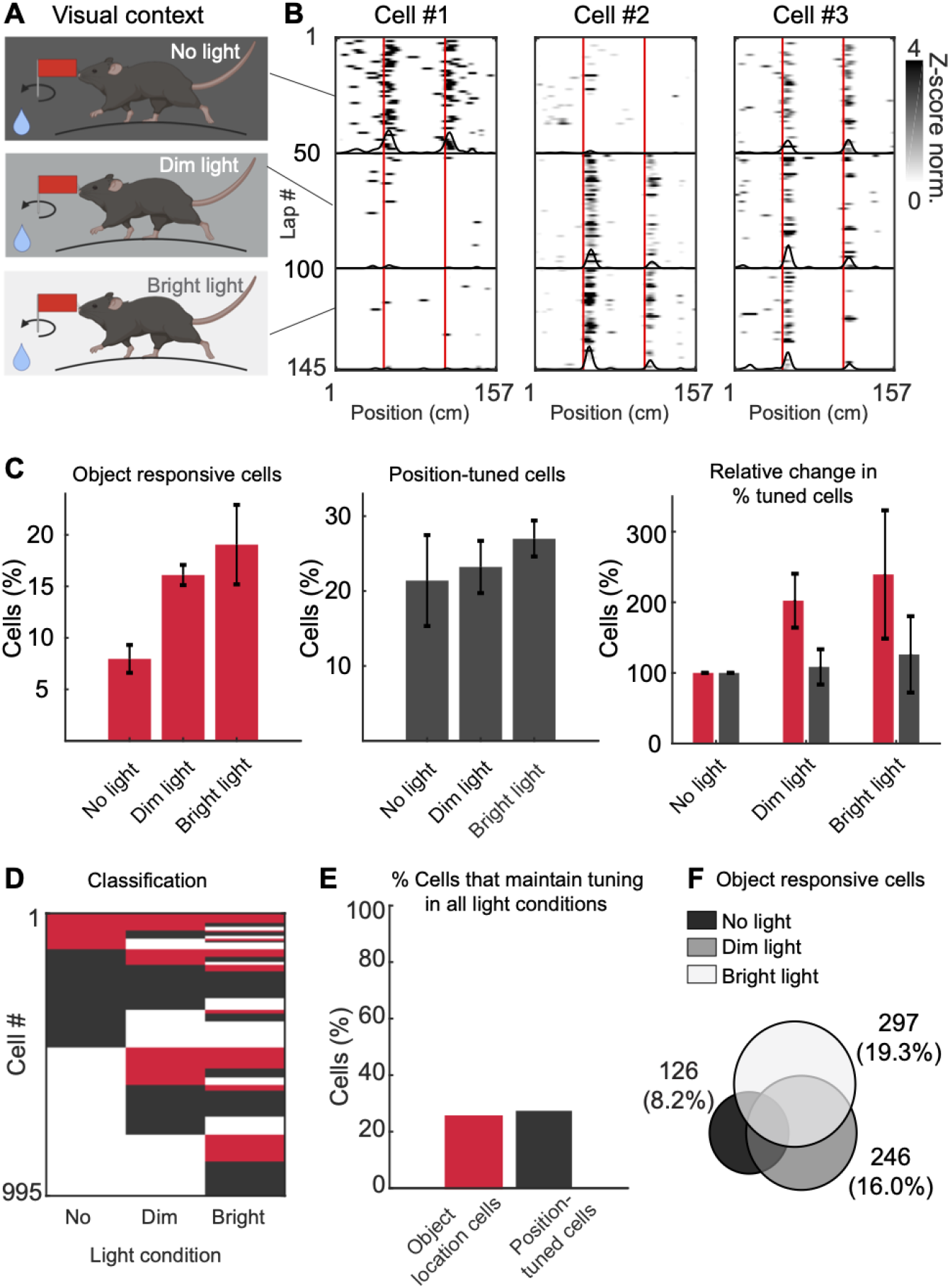
Responses to tactile objects are visual context dependent. (**A**) Mice performed the spatial task while the visual context changed by adjusting the ambient luminance every ~ 50 laps during a session. First, mice ran in complete darkness (“no light”), then under dim ambient light conditions (“dim light”), and finally under bright light conditions (“bright light”). (**B**) Example cells (deconvolved dF/F). Cell #1 only responds in the dark, Cell #2 only when light is present, and Cell #3 responds under all three conditions. (**C**) Percentage of object-responsive cells (left) and position-tuned cells (middle) in each visual context (1510 cells from 5 mice, 5 FOVs; object-responsive cells: “no light” 8.0 ± 1.4 %; “dim light” 16.1 ± 1.0 %; “bright light” 19.1 ± 3.9 %; Position-tuned cells: “no light” 21.4 ± 6.0 %; “dim light” 23.2 ± 3.5 %; “bright light” 27.0 ± 2.4 %). Right: Relative change of classified cells across visual contexts normalized to the task in darkness. (**D**) Single neuron classification across visual contexts (red, object-responsive cell; black, position-tuned cell). Only cells classified in at least one context were included. (**E**) Percentage of object location- and position-tuned cells that keep their tuning identity across all visual contexts (Object location cells 25.4 %; Position-tuned cells 27.4 %). (**F**) Venn diagram of the number of object-responsive cells in each visual context.

Neuronal responses were highly visual-context dependent (Figure 3B). Some neurons responded to tactile objects only in complete darkness, others responded only to congruent visuo-tactile objects, and a minority responded in both darkness and light. Overall, the number of object-responsive neurons almost doubled between dark and bright ambient light (Figure 3C, 1510 neurons, 5 mice, 5 sessions), likely because many RSC neurons are sensitive to the visual features of the moving object. In contrast, the percentage of position-tuned cells did not increase significantly. When we analyzed the tuning properties of object location neurons and position-tuned neurons under different luminance conditions, we observed notable changes (Figure 3D). Most classified cells lost their tuning in ambient light. However, in light, new object-responsive and position-tuned cells emerged. Moreover, object-responsive neurons and position-tuned neurons could switch identities. Overall, only 25.4 % of object location neurons classified in darkness maintained their tuning in light (Figure 3E). Similarly, only 27.4 % of position-tuned neurons classified in darkness maintained their tuning in light. Assessing a Venn diagram of responses under different light conditions showed that neuronal responses to the moving object were highly visual context-dependent, with most neurons responding in bright ambient light (Figure 3F). These data show that both tactile object location neurons and position-tuned neurons in RSC are highly sensitive to changes in visual context.

### RSC develops predictive responses to upcoming tactile objects

When we analyzed the onset time of tactile responses in darkness, we observed that a fraction of cells started to respond before the object was in reach of the whiskers (Figure 4A, reproduced from Figure 1F, and examples in Figure 4B, 8 mice, 10 FOVs, 370 tactile cells out of 3120 cells). We determined the onset times of neuronal responses in relation to the time when the whiskers touch the object based on high-speed videography. We also verified that these early responses were not due to the sound of the motors moving the object (see Methods). Predictive neuronal responses could start up to 50 cm before the whiskers touch the object (Figure 4C). Converted to time, based on the running speed of the animal, these cells could anticipate the object by up to 1 second (Figure 4C). Altogether, 17 % of all object location cells predicted the tactile object (Figure 4D).

**Figure 4.**
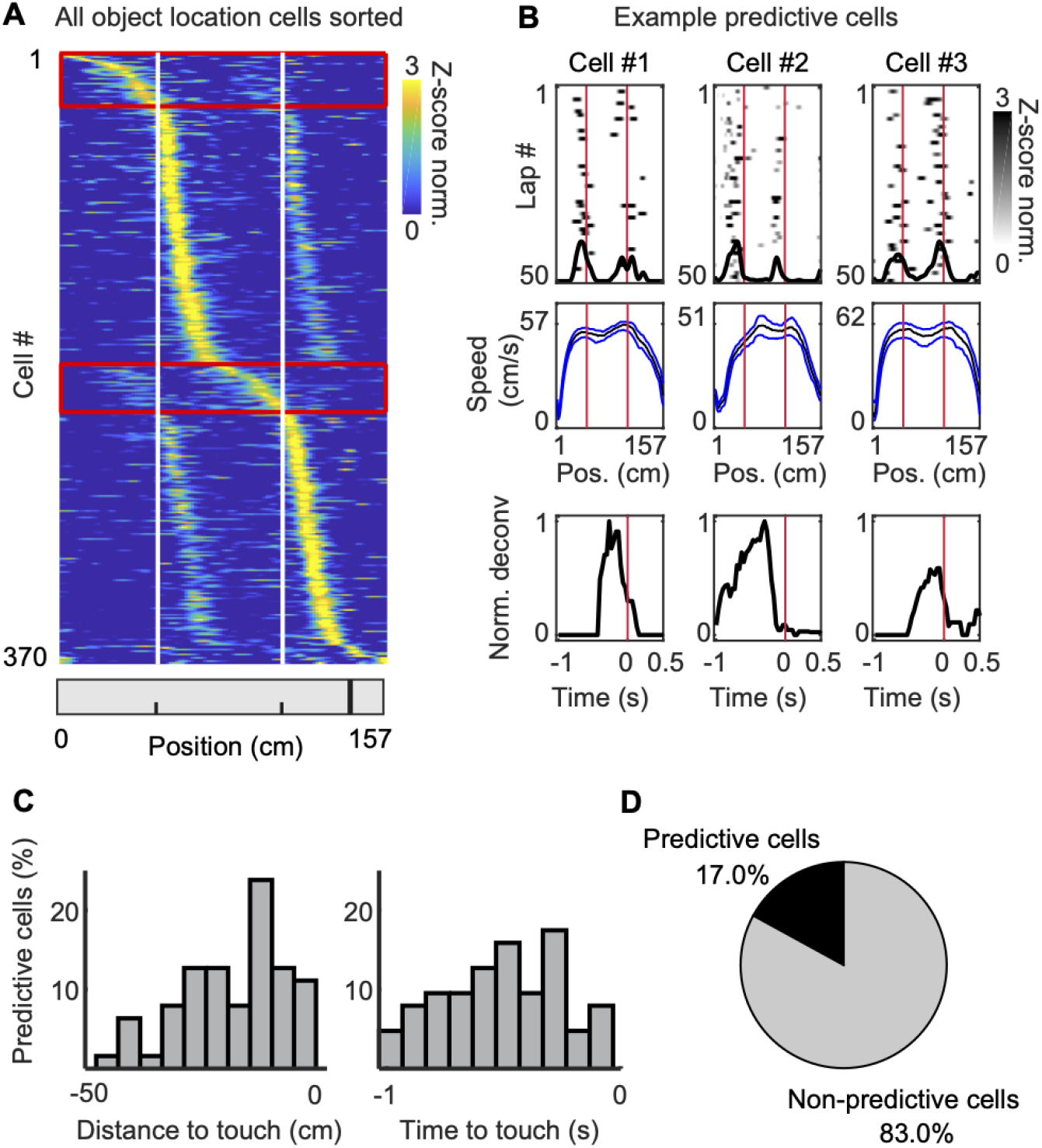
RSC develops predictive responses to upcoming tactile objects. (**A**) Average response of all object location cells, sorted according to the position of peak response (reproduced from Figure 1F). White lines indicate the movement onset of the tactile object. Red boxes indicate cells classified as predictive cells (8 mice, 10 FOVs, 370 object location cells out of 3120 cells). (**B**) Example predictive cells. (Top) Neuronal responses during 50 laps (deconvolved dF/F). Black trace shows the average response. Red lines indicate tactile object movement onset. (Middle) Median running speed. Blue lines indicate the 25th and 75th percentile. (Bottom) Normalized deconvolved dF/F aligned to tactile object onset time. (**C**) Distribution of the response onset of predictive cells aligned to tactile object movement onset, shown on a spatial axis (left) and time axis (right). (**D**) Percentage of predictive- and non-predictive cells among object location neurons.

### Memory traces of tactile object location in RSC

Do tactile objects leave a memory trace in RSC? To test this, we randomly omitted the first tactile object in a subset of laps (Figure 5A, 8 mice, 10 FOVs). After mice ran a minimum of 50 laps, we started omitting the first tactile object randomly in 50 % of laps. For visualization purposes, we sorted the laps according to whether the tactile object was present or omitted (Figure 5A). First, we verified whether omitting a tactile object would change the mouse’s running speed or its estimate of the reward location by analyzing the lick rate, but this was not the case (Figure S2A, B). We found examples of object location cells that lost tuning when the object was omitted (Figure 5B, “non-trace” cell). However, most object location cells responded despite the lacking object (Figure 5B, “trace” cell). When we categorized all object location cells into non-trace and trace cells, we noticed that many trace neurons were active before the tactile object (Figure 5C, bottom traces), indicating that many of the trace cells were also predictive cells (see Figure 4). Indeed, since predictive cells are active before the object, they are agnostic about whether the object will be present or not. Of the object location cells that were classified as predictive neurons, 81% were also classified as trace cells, while only 51% of non-predictive neurons were classified as trace cells (Figure 5D). These data show that most tactile object location cells in RSC maintain a memory trace of where tactile events occurred.

**Figure 5.**
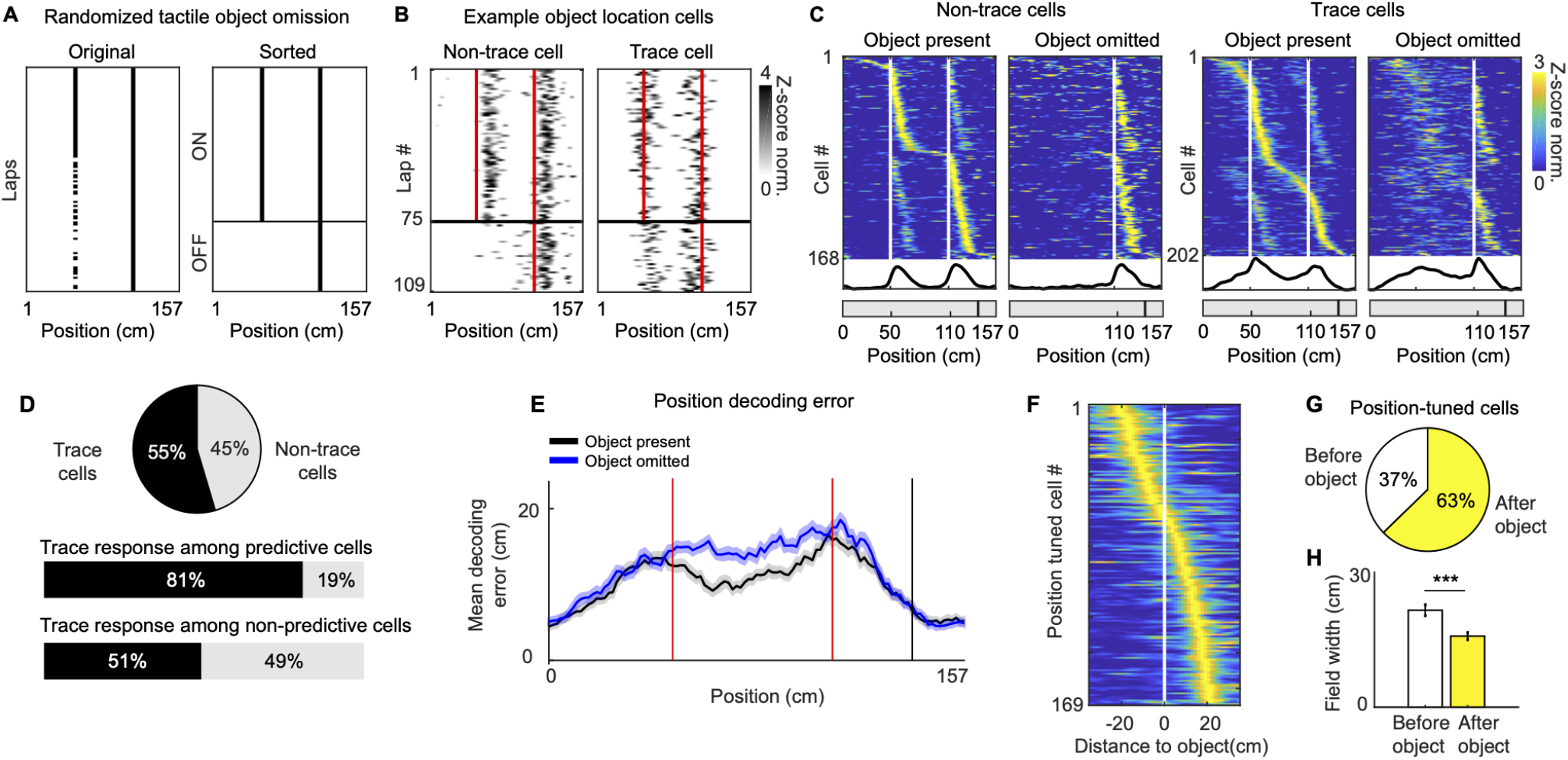
Memory traces of tactile object location in RSC. (**A**) (Left) Half-way a behavior session, the first tactile object was randomly omitted in 50 % of laps. (Right) For visualization, we sorted the laps according to whether the tactile object was present or not. (**B**) Two example object location cells. The non-trace cell example stopped responding when the tactile object was omitted. The trace cell example continued to respond in the absence of the tactile object. Red lines indicate the earliest possible contact time of the whiskers with the object. (**C**) Average response of all trace and non-trace cells, sorted by the position of peak response. The same cells are shown with the first object present (left) or omitted (right). White lines indicate earliest possible contact time of the whiskers with the object. Black line at the bottom shows the average across all cells (8 mice, 10 FOVs, 370 object location cells out of 3120 recorded cells). (**D**) (Top) Percentage of trace and non-trace cells. (Middle) Percentage of trace cells among predictive cells. (Bottom) Percentage of trace cells among non-predictive cells. (**E**) Mean ± SEM decoding error of position using only position-tuned neurons, with the tactile objects present in both locations (control, black), or when the first tactile object is omitted (blue). (**F**) Average response of all position-tuned cells with the peak of their place field at ± 20 cm from either of the two tactile objects. Average responses were sorted by the position of the peak response. (**G**) Percentage of position-tuned cells that were found in the 20 cm window before and after the tactile object. (**H**) Tuning curve width (mean ± SEM) of position-tuned cells in the 20 cm window before or after the tactile object (*** p-value = 12 × 10^−6^, Mann Whitney-U test).

Finally, we tested whether tactile objects influence how well the mouse’s position can be decoded based on the activity of position-tuned neurons. Using Bayesian decoding, we quantified the decoding error as a function of position along the track. This showed that the presence of tactile objects reduced the decoding error (Figure 5E). Further analysis showed that, following contact with an object, there are more position-tuned cells (Figure 5F, G) and the width of the tuning curve of position-tuned cells is significantly smaller (Figure 5H). The combination of these two factors likely decreases the decoding error of spatial position. These data show that tactile events are not only encoded in the activity of object location cells, but also affect the activity of position-tuned neurons and improve the decoding of spatial position.

## Discussion

How do animals use tactile information to detect important objects and remember their location? Here, we asked whether tactile objects are represented in the brain’s spatial network and integrated with the spatial code. We found that neurons in RSC respond to whisker stimulation associated with tactile objects and show several features resembling the properties of a map of the tactile environment. Neuronal responses are highly task and context-dependent, and in well-trained mice, neuronal activity is predictive of upcoming tactile objects. Moreover, neuronal activity persists when tactile objects are removed, indicating that RSC maintains a memory trace of tactile object location.

We found that neurons in RSC encode tactile object location in a task-dependent (Figure 2) and visual context-specific manner (Figure 3). These findings support a framework wherein RSC encodes abstract representations of tactile objects within an environment rather than of tactile objects per se. Several theories propose that the binding of objects to locations occurs in the hippocampus (Connor and Knierim, 2017). However, evidence shows that this may already occur upstream of the hippocampus, in the peri- and postrhinal cortex and entorhinal cortices (reviewed in Connor and Knierim, 2017). There is also evidence that this may happen in the retrosplenial cortex, at least for visual objects (Aggleton and Nelson, 2020; Stacho and Manahan-Vaughan, 2022; Miller et al., 2014; Mitchell et al., 2018; Landeta et al., 2020; Carstensen et al., 2021). Comparing our results to previous investigations is difficult because these were performed in freely moving rodents that explore objects in an arena(Weible et al., 2012; Deshmukh and Knierim, 2013, 2011; Høydal et al., 2019; Tsao et al., 2013; Rivard et al., 2004). There, the animal can inspect and interact with the object using both visual and somatosensory information, which likely includes a mixture of input from the whiskers, limbs, and paws. In such experiments, object-responsive cells may also encode aspects of behaviour related to object exploration and not necessarily object location. Therefore, isolating the contribution of individual sensory modalities requires well-controlled behavioural experiments. Following such an approach, a recent study used head-fixed mice and found that a fraction of hippocampal CA1 cells respond to visuotactile cues attached to a treadmill (Geiller et al., 2017).

These data differed substantially from our RSC data. Hippocampal CA1 responses to cues were not modulated by contextual changes and removing the cues did not cause a lasting memory trace of the cue location. They also found a subset of cells that were active up to 13 cm before the cues, but since experiments were performed in light, they could not exclude that the mouse sees the cues approaching before they are in reach. These findings could indicate key differences between the hippocampus and RSC. Alternatively, the differences may also depend on the task design, as they did not isolate contribution of different sensory modalities and manipulated the contextual cues differently. Therefore, future work should explore how different sensory cues are encoded in the different areas of the brain’s spatial network under standardized behavior conditions.

How do whisker signals reach RSC? There are several possibilities. First, RSC may receive indirect whisker input from the barrel cortex via the posterior parietal cortex (PPC), which forms dense connections with the anterior portion of RSC (Mohan et al., 2019; Merre et al., 2018; Bosman et al., 2011). Second, RSC may receive whisker signals via the same pathways as the hippocampus. Both RSC and the hippocampus receive entorhinal cortex input, although these projections are much sparser in RSC (Sugar et al., 2011; Witter et al., 2017). The entorhinal cortex may receive its tactile information from the barrel cortex via the perirhinal cortex, a key brain area for object recognition (Connor and Knierim, 2017; Ibarra-Casta*ñ*eda et al., 2022; Aronoff et al., 2010). A third possibility is that RSC receives whisker information from the vibrissal motor cortex (Smith and Alloway, 2010) via the claustrum (Brennan et al., 2021). And, finally, there is also an understudied pathway that conveys whisker signals via the spinal trigeminal nucleus to the laterodorsal nucleus of the thalamus (Bosman et al., 2011; Bezdudnaya and Keller, 2008), which in turn, projects to the superficial layers of the agranular RSC (Groen and Wyss, 1992). Of the pathways discussed here, the latter involves the fewest synaptic connections from the mechanoreceptors of the whiskers to RSC.

We found that most object location neurons form a memory trace (Figure 5), and a fraction of these were predictive of upcoming tactile input, in some cases up to 50 cm before the tactile object (Figure 4C). These features resemble some of the properties of recently discovered cells that encode a vectorial signal of the distance and direction between the animal and nearby objects or borders in the environment (Bicanski and Burgess, 2020; Alexander et al., 2020; Wijngaarden et al., 2020). Notably, both vectorial and non-vectoral cells that code for objects and borders can also maintain their response when the object or border is removed, leaving a memory trace of their location (Weible et al., 2012; Deshmukh and Knierim, 2011, 2013; Tsao et al., 2013; O’Keefe, 1976; Poulter et al., 2021). However, because the angle between the animal and object location cannot be assessed when animals run on a linear track, it remains to be determined whether the predictive tactile neurons reported here are vector coding cells.

Interestingly, the neuronal responses that we separated into position-tuned and object location-tuned cells share many similarities. They both lack temporal precision (Figure 1E), they have a similar amplitude and tuning width (Figure 1H), are spatially intermingled (Figure 1J), have a similar sensitivity to visual context (Figure 3D, E) and can even switch identities (Figure 3D). Therefore, these similarities may be based on a common plasticity mechanism. We propose that a prime candidate could be the behavioral timescale plasticity rule (BTSP) that was recently discovered in CA1 pyramidal neurons (Bittner et al., 2017; Zhao et al.,2020). BTSP potentiates only those presynaptic inputs active in a second-long time window around a postsynaptic plateau potential. The essential difference between object location- and position tuned cells could be that they were induced by a plateau potential driven by tactile input and position input, respectively. Notably, using a similar task, we recently reported a preponderance of position-tuned long-range input to RSC from numerous sources (Gianatti et al., 2022). Thus, both tactile-driven and position-driven plateau potentials will lead to a potentiation of position-tuned input. This hypothesis could explain why trace object location cells respond in the absence of an object (Figure 5C), and this could also explain why some cells start to respond up to one second before the tactile object is within reach (Figure 4). Future experiments will have to verify these hypotheses by testing whether RSC neurons can produce plateau potentials and whether this results in object- or position-tuned responses. The head-fixed navigation task with tactile VR proposed here, should be ideally suited to test these ideas and to dissect how tactile objects are encoded and stored in the brain’s spatial network.

## ACKNOWLEDGEMENTS

This work was funded by the European Research Council (ERC Starting Grant #639272) and the Research Council of Norway #274306. We thank Kristin Larsen Sand and Eivind Hennestad for technical assistance. We thank Bruno Pichler (Independent NeuroScience Services; INSS) for developing the custom two-photon microscope and providing technical assistance. Some schematics were created with Biorender.com. The cartoon in Figure 1A is from scidraw.io. Finally, we thank the members of the Vervaeke lab and Matthijs Dorst, Maximiliano Nigro, Shankar Sachidhanandam, Jørgen Sugar, and Jianing Yu for providing critical comments that significantly improved a draft of the manuscript.

## AUTHOR CONTRIBUTIONS

AL and KV designed the study. AL performed all experiments and analysis. AL and KV wrote the paper. All authors discussed the results.

## Methods

### Resource availability

#### Lead contact

Further information and resource requests should be directed to and will be fulfilled by the lead contact Koen Vervaeke.

#### Materials availability

This study did not generate new unique reagents.

#### Data and code availability

- Ca^2+^ imaging and tracing data reported in this paper will be shared by the lead contact upon request.
- All original code will be deposited on GitHub and is publicly available as of the date of publication. Any additional information required to reanalyze the data reported in this paper is available from the lead contact upon request.

### Experimental model and subject details

We used male and female Thy1-GCaMP6s mice (GP 4.3 line #024275 from JAX, (Dana et al., 2014)) between 3 – 8 months old at the time of surgery. We housed mice in groups with 2-4 littermates on a reversed 12-hour light/ 12-hour dark cycle, and we carried out experiments during their dark phase. Mice were kept in a humidity and temperature-controlled environment. Mice received water drop rewards throughout the task, and we kept them on a water restriction regime as described in (Guo et al., 2014). All procedures were approved by the Norwegian Food Safety Authority (project: FOTS 6590, 7480, 19129) and experiments were performed in accordance with the Norwegian Animal Welfare Act.

### Methods details

#### Surgery

Surgery was carried out under isoflurane anesthesia (3 % induction, 1 % maintenance) while maintaining body temperature at 37°C with a heating pad (Harvard Apparatus). We delivered a subcutaneous injection of 0.1 mL Marcaine (bupivacaine 0.25 % m/V in sterile water) at the scalp incision site and administered post-operative analgesia (Temgesic, 0.1 mg/kg) subcutaneously. Several weeks before experiments began, we implanted the head bar and cranial window. The center of the window was −2.2 mm AP from bregma. The window consisted of a circular outer window (3.5 mm diameter, #1.5 coverslip glass) affixed to a circular inner window (2.5 mm diameter, #2 coverslip glass) with optical adhesive (Norland Optical Adhesive; ThorLabs NOA61). We held the window in place by applying gentle pressure so that the outer window would fit into the thinned skull area and flush with the skull’s surface. We used heated agar (1 %; Sigma #A6877) to seal any open spaces between the skull and edges of the glass window, and we affixed the window to the skull with cyanoacrylate glue. A detailed protocol of the surgery can be found in (Holtmaat et al., 2009).

#### Water restriction

Starting a minimum 1 week after surgery, we placed mice on water restriction (Guo et al., 2014). Mice received 1-1.5 ml of water per day while we monitored their body weight to ensure they maintained more than 80% of the initial body weight.

#### Tactile VR setup

Mice were trained to find the location of a water reward on a 50 cm diameter Styrofoam wheel (circumference 157 cm). On the left side of the animal, we positioned a rotary servo motor (Zaber NM08AS-T4; controlled by Zaber X-MCB2 24V motor controller) and to the shaft we attached a 2 × 6 cm piece of cardboard. The distance of the motor was adjusted such that the cardboard brushed the outer one-third of the whiskers. The rotation speed of the rotary motor was controlled closed loop by the running speed of the mouse which was recorded using an optical rotary encoder attached to the running wheel axis. The coupling gain was adjusted so the cardboard brushed the whiskers at the same speed as the mouse’s running speed, simulation the sensation of running passed an object. The rotary motor was triggered to start moving at positions 45 cm and 105 cm along the track, relative to the location of the water reward. It takes about 5 cm of forward movement by the mouse before the motors rotate the object in reach of the whiskers. Therefore, the tactile object reached the whiskers at positions 50 cm and 110 cm. Once the mouse passed the tactile object, the motor was reset to its home position, out of reach of the whiskers, until the next tactile object.

The behavior was automated used custom routines in LabView (2018, National Instruments) to control all actuators, sensors, data acquisition, to record behavioral parameters, and to trigger the microscope to start and stop recording. All behavioral parameters were recorded at 5 kHz. The absolute position on the running wheel was based on the rotary encoder and additionally calibrated using an IR emitter-receiver pair (transceiver) facing the running wheel. A small black cue passing the transceiver then signaled each time a full lap passed. In addition to the motor-controlled tactile object, a strip of fine sandpaper (2 cm wide) was attached to the running wheel 17 cm before the reward location. The behavior setup was positioned under a custombuild two-photon microscope in a light-tight enclosure built from Thorlabs parts (25 mm optical rails, Thorlabs, XE25L48), blackout hardboard (Thorlabs, TB4) and blackout fabric (Thorlabs, BK5). It was key that no light could enter the enclosure and additionally, the room lights where the microscope was located were turned off.

#### Passive whisker stimulation

In a subset of experiments (Figure 2), we passively stimulated the whiskers. The running wheel was clamped to immobilize the mouse and at random intervals the motor brushed the tactile object over the whiskers at a speed matching the running speed of the mouse in a normal session, typically 40-50 cm/s. After about 15 trials, the wheel was unlocked allowing the mouse to freely run again and the passive stimulation was again repeated during running for about 15 laps. The stimulations were separated by at least 5 seconds.

#### Dark/Light experiments

In experiments comparing responses in darkness and two different levels of ambient light, we used a Thorlabs white LED (MNWHL4) to illuminate the inside of the enclosure. The LED was positioned 0.5 meter behind the mouse and controlled by a Thorlabs LED driver (LEDD1B) which was modulated through a 5V input drive via LabView. The current applied to the LED was: no-light: 0 mA; dim light: 7.2 mA; bright light: 24 mA. The mice were trained on the task in darkness before the dark/light experiments were performed.

#### Whisker imaging and pupil tracking

High speed imaging of the whiskers was done using a NORPIX imaging system with a Basler Aca2000-340kmNIR camera and Streampix-6 acquisition software at 200 Hz. To illuminate the whiskers, we used a 940 nm IR LED (Thor Labs M940L3). Whisker imaging was synchronized with two-photon imaging using the microscope frame clock to send TTLs to a small IR LED positioned in the whisker camera’s field of view (see Video S1). This TTL was set to 5 V during the first half of a microscope frame and 0 V for the second half. A custom MATLAB script then searched the whisker imaging frames, and every frame where the LED switched on was matched up with the corresponding two-photon imaging frame. In a subset of mice, we performed pupil tracking to look for correlations in pupil parameters such as diameter, and pupil movements in the yaw and pitch rotation axis (Figure S1). This was done using the same camera and software for whisker imaging, but at 50 Hz. Extraction of pupil parameters was done using custom code in MATLAB.

#### Behavioral training

Mice were water restricted for 7-10 days before their first exposure to the training environment (Guo et al.,2014). To habituate the animals to being head-fixed and walk on the Styrofoam wheel, the mice were exposed gradually to the setup, typically only 3-5 minutes on the first and second day of training. As mice got more confident, the duration was increased to the point where they ran typically 300 meters during a 20-minute session. When mice were running comfortably for a minimum of 80 meters with a parabolic running profile (see Figure 1C), the motor-controlled tactile objects were introduced. Thereafter, the mice were trained to run with the tactile objects present for at least a week before imaging experiments started.

#### Two-photon imaging

Imaging was performed on a custom-built two-photon microscope designed to provide a large space under the objective, allowing the large Styrofoam running wheel of the VR setup (Hennestad et al., 2021). The microscope was built in collaboration with Independent Neuroscience Services (INSS). All imaging data was acquired at 31 fps (512 × 512 pixels) using the open-source acquisition software SciScan (written in LabVIEW). The excitation wavelength was 950 nm using a MaiTai DeepSee ePH DS laser (Spectra-Physics), where the average power measured under the objective (N16XLWD-PF, Nikon) was typically 60-110 mW. The field of view width varied between 500-600 *μm*. Photons were detected using GaAsP photomultiplier tubes (PMT2101/M, Thorlabs). The primary dichroic mirror was a 700 nm LP (Chroma), and the photon detection path consisted of a 680 nm SP filter (Chroma), a 566 nm LP dichroic mirror (Chroma), and a 510/80 nm BP filter (Chroma). The field of view center for recordings was between −1.7 and −3.2 mm anterior-posterior, and 0.3 to 0.7 mm medial-lateral, relative to bregma. Imaging depth varied between 100-220 *μm* below the cortical surface, corresponding to Layer 2/3 in the agranular RSC.

#### Motion correction, image segmentation and signal extraction

Images were registered using a combination of custom and published code. Images were first de-stretched to correct for distortions resulting from the sinusoidal speed profile of the resonance scan mirror of the microscope. Next, both rigid and non-rigid motion correction was performed using custom written scripts utilizing NoRMCorre (Pnevmatikakis et al., 2016). Registered images were then loaded into a custom written MATLAB GUI to manually draw regions of interest (ROIs) around neuronal somas. When ROIs were overlapping the overlapping region was excluded. A doughnut-shaped ROI of surrounding neuropil was automatically created for each ROI, by dilating the ROI so that the doughnut area was four times larger than the soma ROI area. If this doughnut overlapped with another soma ROI, then that soma ROI was excluded from the doughnut.

The ROI fluorescence is computed as a fractional change dF/F according to *f*(*t*) = (*F*(*t*) − *F*_0_)/*F*_0_, with *F*_0_ being the baseline defined as the 20th percentile of the signal. This was calculated for both the soma and neuropil ROI. Then the neuropil dF/F was subtracted from the soma dF/F, and finally, a correction factor was added to ensure that the soma dF/F remained positive. The dF/F for each cell was then deconvolved using the CaImAn package (Giovannucci et al., 2019). The resulting deconvolved signals are given in arbitrary units (a.u.).

## QUANTIFICATION AND STATISTICAL ANALYSIS

Time series for dF/F, deconvolved dF/F and behavioral data were processed and analyzed using Matlab. The behavioral time series were first binned to match the two-photon microscope frame rate (31 Hz) by finding the behavioral data sample closest in time to the imaging frame.

### Running speed analysis

The running speed was computed by calculating the change in distance between two samples. Using the wheel circumference (**2** · *π* · *radius* of 25 cm) and dividing it by the number of ticks in one full rotation of the optical encoder (2000 ticks), an estimated running speed was obtained. The running speed was then smoothed using a 0.1 second moving average to remove unwanted spikes. For each sample point, the absolute position on the wheel was found by computing the modulo of the number of ticks T(t) by max number of ticks for one round:

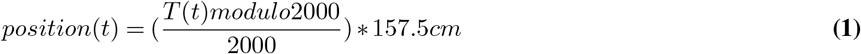

The position was then offset by the position of the first sample where the wheel position LED diode was triggered, which was set to be position 0 cm. The lap number at sample t was computed by the following:

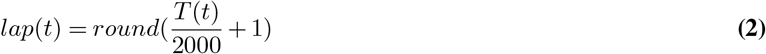

where T(t) is the tick count at sample t. Both position and lap time series were down sampled to match the imaging frames by computing the mean value at each frame. Finally, the position along the track was binned into 1.5 cm bins, creating 105 bins along the linear track.

### Tactile object movement detection

The movement of the motor-controlled tactile objects was detected for each lap either by using the whisker imaging data, or by recording the rotary motor speed signal. For all plots where a line is used to visualize the object position, and for analysis that required the object position, we used bin 32 and 72 (48 cm and 108 cm), which was found to be the last bin where no whisker-object interaction was observed.

### Average tuning curves

For each cell, we created an average tuning curve to quantify the response profile along the linear track. First, the linear track was divided into 1.5 cm bins, giving 105 bins. An occupancy count was created by counting the number of time bins the animal had been within each of the 105 bins. For each cell the deconvolved dF/F response at each bin was summed and later divided by the occupancy count in each bin, creating the final occupancy normalized averaged tuning curve.

### Classification of position-tuned cells

Classification was similar to (Mao et al., 2017). Position tuned cells were detected by finding significant peak responses in the averaged tuning curve. Peaks were detected using the MATLAB Signal Processing Toolbox function findpeaks(). Each peak that was detected had to be separated by a minimum distance of 40 cm to the surrounding peaks, have a minimum width of 2 cm and a maximum width of 120 cm. A maximum of 3 peaks could be detected by the algorithm. A “response threshold” was used to detect a peak, which was defined as 30% of the difference between the maximum and minimum of the average tuning curve. Because the linear track is circular, meaning position 0 cm is the same as position 157 cm, the first 45 cm of the position map was added to the end of the position map when detecting peaks to allow the full shape of the response peaks to be present. Finally, the neural response had to cross the response threshold within its tuning field in 1/3 of laps. To find the field width of position-tuned cells, we looked forward and backwards at each detected peak to find the positions where the response would fall below the “response threshold” and set the field width to be the number of bins around the peak where the cell showed continuous response above this threshold.

### Classification of object location cells

First, we detected cells that, based on their tuning curve, showed a significant response ± 15 cm from the tactile object position. For classification, the tuning curve was first smoothed using a Gaussian filter with sigma = 30 cm. A cell was then classified if it had a response peak at both positions where the tactile object was presented. The response peaks in the averaged tuning curve were found using the MATLAB Signal Processing Toolbox function findpeaks(), set to search a max number of 4 peaks. The peaks were limited to a max width of 75 cm, and a minimum distance between peaks of 30 cm. A peak was deemed significant if (1) it was greater than 25% of the difference between the max and min of the average tuning curve and (2) it was higher than the reliability criteria. The reliability criteria required that a response was present on a minimum of 20% of the laps for the most prominent of the two peaks, or a minimum of 10% of the laps for the least prominent of the two peaks.

### Classification of trace and non-trace object location cells

A trace cell showed a significant response at the position of the missing tactile object. To determine this, we first combined all laps where the tactile object was omitted and created an average tuning curve. The baseline activity of each tactile cell was computed by taking the 10 least active bins of the average tuning curve and calculating the mean and standard deviation of these bins. A tactile cell was classified as a trace tactile cell if (1) the maximum response at ± 15 cm from the tactile object crossed a threshold set as the baseline mean + (baseline standard deviation * 3) and (2) this threshold was crossed in at least 20% of laps. Non-trace cells, trace cells and position tuned cells were mutually exclusive cell classes.

### Passive stimulus responding cells

For a given cell, dF/F responses were collected in a window of 4 seconds before and 4 seconds after the stimulation, for each trial. This was done for both immobile passive stimulation and for passive stimulation during running, resulting in two matrices of n trials by 8 seconds. We then classified responses for each matrix using shuffling by randomly shifting the response in each trial (using MATLABs function circshift()). An average response was computed for the original matrix and for the shuffled matrix. We created 1000 shuffled matrices and computed the average response of each. A response was deemed significant if the average response in a 650 ms window starting 350 ms after the stimulation was larger than the 95th percentile of the shuffled responses in the same time window.

### Predictive cells

The average tuning curve of all object location cells was used to find predictive cells. All cells that had their maximum response along the track before the first tactile object onset and all cells that had a maximum response up to 35 cm before the second tactile object was deemed to be predictive. To find the onset position of each predictive cell, we found the earliest position along the track that the cell showed an average response above the 95th percentile of the cells’ average tuning curve. To find the *onset time* of each predictive cell, we first found each time bin where the mouse entered the tactile object at the first object position and created a matrix containing the deconvolved dF/F signal in a time window of 1 second before. We then averaged this time aligned response and used it to find the time bin where the signal first crossed the 95th percentile of the averaged deconvolved dF/F signal.

### Light responsive cells

For experiments in light, cells were classified as object location cells as described above. The experiments had three levels of light; no light, dim light, and bright light. Cells were tested for being responsive in each visual context by using the laps for each of these light conditions.

### Analysis of changes in tuning across light contexts

Cells were classified as position-tuned or object location cells across the three light contexts as detailed above. To compute similarity across contexts, we computed the percentage of cells that stayed within their class across all three contexts.

### Population decoding

We used a Bayesian algorithm (Zhang et al., 1998) as a decoding model (Mao et al., 2017) to estimate the position of the animal based on the activity of all position-tuned cells. This was done to compare the spatial information in laps where the tactile object was present or omitted. Data from 10 session (8 mice) was included. Decoding was done only for periods where the running speed of the animal was above 1 cm/s, using the deconvolved dF/F traces, excluding all cells that had no deconvolved transients. Deconvolved signals were smoothed using a Gaussian filter with sigma = 5 cm. The time series were later binned (0.19 s) using non-overlapping time bins selected to minimize decoding error. The data was split into laps where the tactile object was present or omitted. Training data was generated using odd laps where the tactile object was present. Two sets of test data were used, even laps when the object was presented (control), or all laps when the object was not presented (omitted). Mean decoding error and SEM are reported by combining the individual laps across all included sessions. The decoding error was computed as the absolute difference between the actual and decoded position of the animal at each sample bin. The decoded position was selected from the maximum likelihood given the activity of all position-tuned cells within a time bin. The maximum likelihood was found by computing the probability of the animal’s position at each position given the population activity:

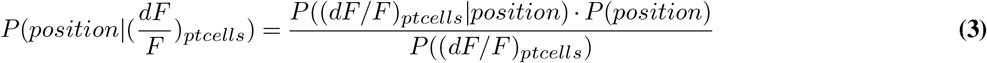

Where *position* is the binned position at the given time bin, (*dF/F*)_*ptcells*_ is the population activity of all position-tuned cells, and P is probability. P(*position*) was computed using the occupancy in each position bin.

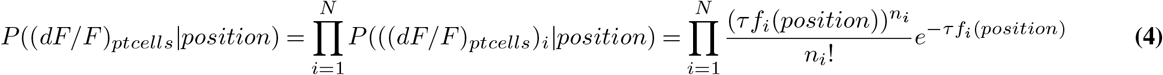

where *τ* is the time bin (0.19 s), *f_i_*(*position*) is the computed position tuning curve for the i-th neurons, N is the total number of neurons; and *n_i_* is the mean activity of the i-th neuron within that time bin. This gives:

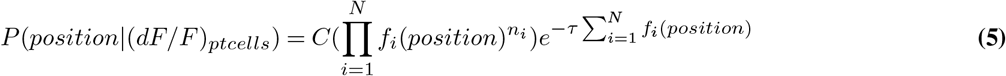

where C serves as a normalization factor so that *P*(*position*|(*dF/F*)_*ptcells*_) sums to 1.

### Analyzing position-tuned cells around tactile objects

To find position-tuned cells that were active around the tactile objects, we first quantified cells that were position-tuned using the first 50 laps where the tactile object was presented. For each of these cells we created an average tuning response and picked only cells with a single firing field along the track for further analysis. We subsequently selected cells that had their peak activity ± 21 cm from either the first or the second tactile object. Field width of each position-tuned cells were found as described under “*Classification of position tuned cells*”.

### Eye movement analysis

The pupil was back-illuminated by the IR laser light used for two-photon imaging. First, a raster map was produced for each of the eye signals; the horizontal (yaw) eye movement, the vertical (pitch) eye movement, and pupil diameter. Because experiments were performed in the dark, the pupil was fully dilated. Therefore, reductions in eye diameter (Figure S1D, E) reflect eyeblinks rather than pupil contractions. To determine correlations between the eye blinks and neuronal activity, we resampled the 50 Hz pupil video to match the two-photon imaging capturing at 31 Hz. An eye blink event was defined as a drop in eye radius below a baseline value. The baseline was set to be the mean - 2 × the standard deviation of eye radius in a window of 30 cm before the tactile object was presented, up to the point of object presentation. To test which neurons were significantly correlated with eye squints, we performed Wilcoxon rank sum test on each recorded neuron. To do so, we found the max response of each neuron in a 15 cm window after the tactile object was presented, and split the data into two vectors, one containing the responses during eye blinks, and one containing the responses when there were no eye blinks. The rank sum test used alpha = 0.05 and a left tail.

## Supplementary Figures

**Figure S1.**
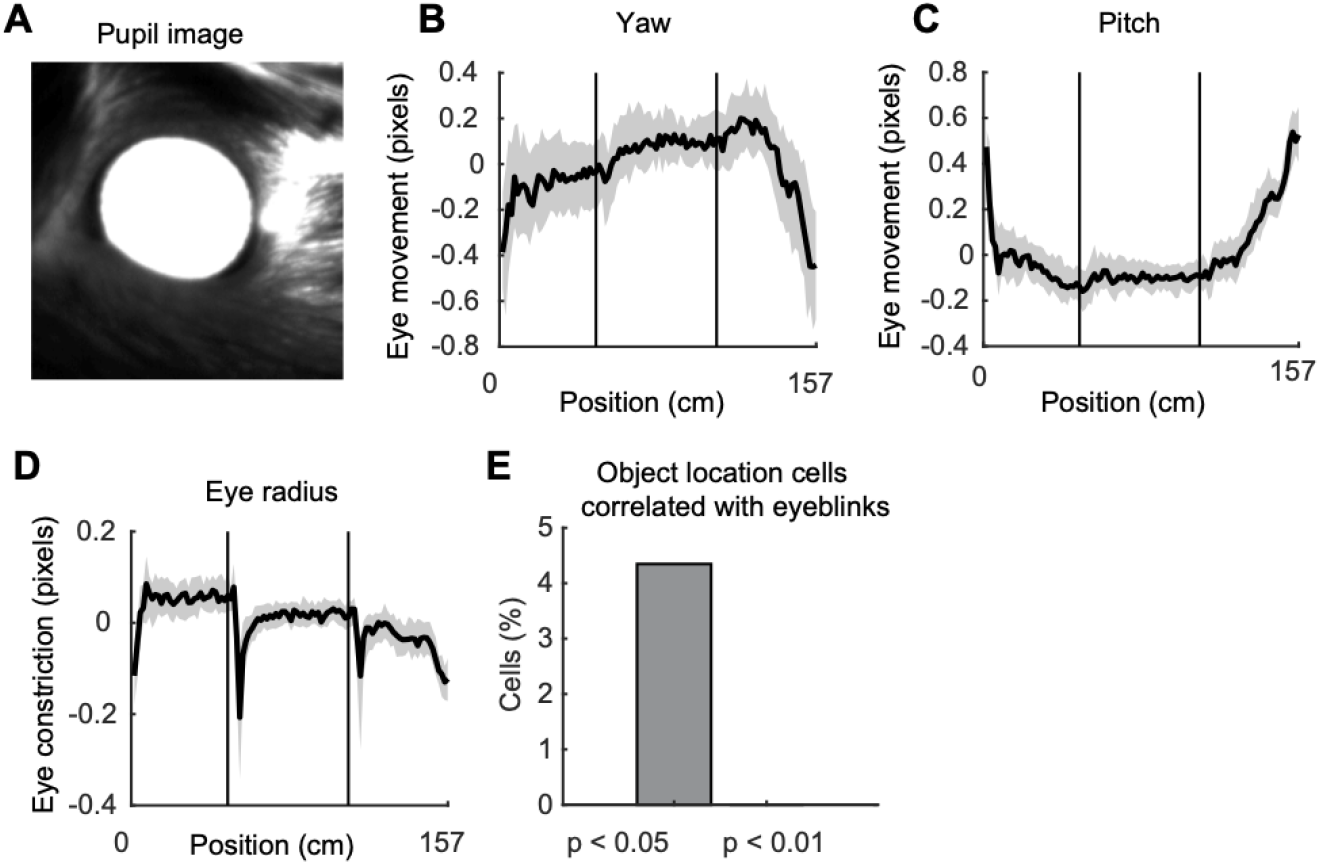
Effects of tactile stimuli on the eye pupil.. (**A**) Example image of the mouse pupil. The pupil is back-illuminated by the two-photon IR laser light used during neuronal recordings. (**B**-**C**) Median movement of the pupil in the yaw (B) and pitch (C) eye rotation axis. Shaded area shows 25th and 75th percentile. Black lines indicate position of the tactile object movement onset (3 mice, 3 sessions). (**D**) Pupil diameter changes due to brief eye squint events following tactile object contact. (**E**) Percentage of object location cells significantly correlated with brief eye squint events.

**Figure S2.**
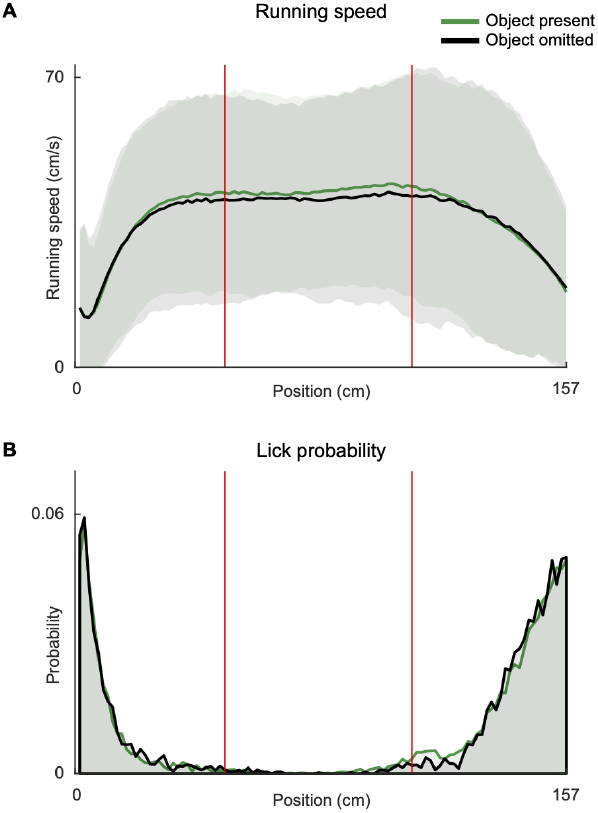
Running speed and lick rate with and without tactile object. (**A**) Average running speed of mice across all sessions, with tactile objects present (black) and with the first tactile object omitted (red) (8 mice, 10 sessions). Shaded error bar indicates two standard deviations from the mean. Red lines indicate tactile object position. (**B**) Lick probability averaged across all mice.

## Supplementary Video Captions

**Video S1. Tactile responses in RSC during a spatial task in darkness.** Example experiment showing videography of the whisker and tactile object movements (top), and the responses of an example object location cell during 18 laps (middle panels). Blue ticks indicate deconvolved dF/F events. Bottom, neuronal response of the example cell (dF/F, black) and deconvolved events (blue). Red lines indicate tactile events.

## Bibliography

Aggleton, J. P. and Nelson, A. J. (2020). Distributed interactive brain circuits for object-in-place memory: A place for time? Brain and Neuroscience Advances.

Alexander, A. S., Carstensen, L. C., Hinman, J. R., Raudies, F., Chapman, G. W., and Hasselmo, M. E. (2020). Egocentric boundary vector tuning of the retros-plenial cortex. Science Advances, (8).

Alexander, A. S. and Nitz, D. A. (2015). Retrosplenial cortex maps the conjunction of internal and external spaces. Nature Neuroscience, 18(8):1143–1151.

Aronoff, R., Matyas, F., Mateo, C., Ciron, C., Schneider, B., and Petersen, C. C. (2010). Long-range connectivity of mouse primary somatosensory barrel cortex. European Journal of Neuroscience, 31(12):2221–2233.

Ayaz, A., Stäuble, A., Hamada, M., Wulf, M.-A., Saleem, A. B., and Helmchen, F. (2019). Layer-specific integration of locomotion and sensory information in mouse barrel cortex. Nature Communications, 10(1):2585.

Bar, M. (2004). Visual objects in context. Nature reviews. Neuroscience, 5(8):617–629.

Bellistri, E., Aguilar, J., Brotons-Mas, J. R., Foffani, G., and Prida, L. M. d. l. (2013). Basic properties of somatosensory-evoked responses in the dorsal hippocampus of the rat. The Journal of Physiology, 591(10):2667–2686.

Bezdudnaya, T. and Keller, A. (2008). Laterodorsal nucleus of the thalamus: A processor of somatosensory inputs. Journal of Comparative Neurology, 507(6):1979–1989.

Bicanski, A. and Burgess, N. (2020). Neuronal vector coding in spatial cognition. Nature Reviews Neuroscience, 21(9):453–470.

Bittner, K. C., Milstein, A. D., Grienberger, C., Romani, S., and Magee, J. C. (2017). Behavioral time scale synaptic plasticity underlies CA1 place fields. Science, 357(6355):1033–1036.

Bosman, L. W. J., Houweling, A. R., Owens, C. B., Tanke, N., Shevchouk, O. T., Rahmati, N., Teunissen, W. H. T., Ju, C., Gong, W., Koekkoek, S. K. E., and Zeeuw, C. I. D. (2011). Anatomical Pathways Involved in Generating and Sensing Rhythmic Whisker Movements. Frontiers in Integrative Neuroscience, 5:53.

Brennan, E. K., Jedrasiak-Cape, I., Kailasa, S., Rice, S. P., Sudhakar, S. K., and Ahmed, O. J. (2021). Thalamus and claustrum control parallel layer 1 circuits in retrosplenial cortex. eLife.

Carstensen, L. C., Alexander, A. S., Chapman, G. W., Lee, A. J., and Hasselmo, M. E. (2021). Neural responses in retrosplenial cortex associated with environmental alterations. iScience, 24(11):103377.

Chen, L. L., Lin, L. H., Green, E. J., Barnes, C. a., and McNaughton, B. L. (1994). Head-direction cells in the rat posterior cortex. I. Anatomical distribution and behavioral modulation. Experimental Brain Research, 101(1):8–23.

Cheung, J. A., Maire, P., Kim, J., Lee, K., Flynn, G., and Hires, S. A. (2020). Independent representations of self-motion and object location in barrel cortex output. PLoS Biology, (11).

Claessen, M. H. and Ham, I. J. v. d. (2017). Classification of navigation impairment: A systematic review of neuropsychological case studies. Neuroscience & Biobehavioral Reviews, 73:81–97.

Connor, C. E. and Knierim, J. J. (2017). Integration of objects and space in perception and memory. Nature neuroscience, 20(11):1493–1503.

Couto, J., Kandler, S., Mao, D., McNaughton, B., Arckens, L., and Bonin, V. (2019). Spatially segregated responses to visuo-tactile stimuli in mouse neocortex during active sensation. bioRxiv.

Cowansage, K. K., Shuman, T., Dillingham, B. C., Chang, A., Golshani, P., and Mayford, M. (2014). Direct reactivation of a coherent neocortical memory of context. Neuron, 84(2):432–441.

Czajkowski, R., Jayaprakash, B., Wiltgen, B., Rogerson, T., Guzman-Karlsson, M. C., Barth, A. L., Trachtenberg, J. T., and Silva, A. J. (2014). Encoding and storage of spatial information in the retrosplenial cortex. Proceedings of the National Academy of Sciences, 111(23):8661–8666.

Dana, H., Chen, T.-W., Hu, A., Shields, B. C., Guo, C., Looger, L. L., Kim, D. S., and Svoboda, K. (2014). Thy1-GCaMP6 Transgenic Mice for Neuronal Population Imaging In Vivo. PLoS ONE, 9(9):e108697.

Deshmukh, S. S. and Knierim, J. J. (2011). Representation of Non-Spatial and Spatial Information in the Lateral Entorhinal Cortex. Frontiers in Behavioral Neuroscience, 5:69.

Deshmukh, S. S. and Knierim, J. J. (2013). Influence of local objects on hippocampal representations: Landmark vectors and memory. Hippocampus, 23(4):253–267.

Diamond, M. E., Heimendahl, M. v., Knutsen, P. M., Kleinfeld, D., and Ahissar, E. (2008). ‘Where’ and ‘what’ in the whisker sensorimotor system. Nature Reviews Neuroscience, 9(8):601–612.

Fischer, L. F., Soto-Albors, R. M., Buck, F., and Harnett, M. T. (2020). Representation of visual landmarks in retrosplenial cortex. eLife, 9:e51458.

Franco, L. M. and Goard, M. J. (2021). A distributed circuit for associating environmental context with motor choice in retrosplenial cortex. Science Advances, 7(35):eabf9815.

Fyhn, M., Hafting, T., Treves, A., Moser, M.-B., and Moser, E. I. (2007). Hippocampal remapping and grid realignment in entorhinal cortex. Nature, 446(7132):190–194.

Gallace, A. and Spence, C. (2014). Chapter 5-A memory for touch. pages 111–146.

Gallero-Salas, Y., Han, S., Sych, Y., Voigt, F. F., Laurenczy, B., Gilad, A., and Helmchen, F. (2021). Sensory and Behavioral Components of Neocortical Signal Flow in Discrimination Tasks with Short-Term Memory. Neuron, 109(1):135–148.e6.

Geiller, T., Fattahi, M., Choi, J.-S., and Royer, S. (2017). Place cells are more strongly tied to landmarks in deep than in superficial CA1. Nature Communications, 8(1):14531.

Gener, T., Perez-Mendez, L., and Sanchez-Vives, M. V. (2013). Tactile modulation of hippocampal place fields. Hippocampus, 23(12):1453–1462.

Gianatti, M., Garvert, A. C., and Vervaeke, K. (2022). Diverse long-range projections convey position information to the retrosplenial cortex. bioRxiv, page 2022.09.18.508427.

Giovannucci, A., Friedrich, J., Gunn, P., Kalfon, J., Brown, B. L., Koay, S. A., Taxidis, J., Najafi, F., Gauthier, J. L., Zhou, P., Khakh, B. S., Tank, D. W., Chklovskii, D. B., and Pnevmatikakis, E. A. (2019). CaImAn an open source tool for scalable calcium imaging data analysis. eLife, 8:e38173.

Groen, T. v. and Wyss, J. M. (1992). Connections of the retrosplenial dysgranular cortex in the rat. The Journal of comparative neurology, 315(2):200–216.

Guo, Z. V., Hires, S. A., Li, N., O’Connor, D. H., Komiyama, T., Ophir, E., Huber, D., Bonardi, C., Morandell, K., Gutnisky, D., Peron, S., Xu, N.-l., Cox, J., and Svoboda, K. (2014). Procedures for Behavioral Experiments in Head-Fixed Mice. PLoS ONE, 9(2):e88678.

Harvey, C. D., Coen, P., and Tank, D. W. (2012). Choice-specific sequences in parietal cortex during a virtual-navigation decision task. Nature, 484(7392):62–68.

Hennestad, E., Witoelar, A., Chambers, A. R., and Vervaeke, K. (2021). Mapping vestibular and visual contributions to angular head velocity tuning in the cortex. Cell Reports, 37(12):110134.

Holtmaat, A., Bonhoeffer, T., Chow, D. K., Chuckowree, J., Paola, V. D., Hofer, S. B., Hübener, M., Keck, T., Knott, G., Lee, W.-C. A., Mostany, R., Mrsic-Flogel, T. D., Nedivi, E., Portera-Cailliau, C., Svoboda, K., Trachtenberg, J. T., and Wilbrecht, L. (2009). Long-term, high-resolution imaging in the mouse neocortex through a chronic cranial window. Nature Protocols, 4(8):1128–1144.

Høydal, Ø. A., Skytøen, E. R., Andersson, S. O., Moser, M.-B., and Moser, E. I. (2019). Object-vector coding in the medial entorhinal cortex. Nature, 568(7752):400–404.

Ibarra-Castañeda, N., Moy-Lopez, N. A., and González-Pérez, O. (2022). Tactile information from the vibrissal system modulates hippocampal functioning. Current Research in Neurobiology, 3:100034.

Itskov, P. M., Vinnik, E., and Diamond, M. E. (2011). Hippocampal Representation of Touch-Guided Behavior in Rats: Persistent and Independent Traces of Stimulus and Reward Location. PLoS ONE, 6(1):e16462.

Kentros, C. G., Agnihotri, N. T., Streater, S., Hawkins, R. D., and Kandel, E. R. (2004). Increased attention to spatial context increases both place field stability and spatial memory. Neuron, 42(2):283–295.

Knutsen, P. M. and Ahissar, E. (2009). Orthogonal coding of object location. Trends in Neurosciences, 32(2):101–109.

Landeta, A. B. d., Pereyra, M., Medina, J. H., and Katche, C. (2020). Anterior ret-rosplenial cortex is required for long-term object recognition memory. Scientific Reports, 10(1):4002.

Maguire, E. (2001). The retrosplenial contribution to human navigation: A review of lesion and neuroimaging findings. Scandinavian Journal of Psychology, 42(3):225–238.

Mao, D., Kandler, S., McNaughton, B. L., and Bonin, V. (2017). Sparse orthogonal population representation of spatial context in the retrosplenial cortex. Nature communications, pages 1–9.

Mao, D., Neumann, A. R., Sun, J., Bonin, V., Mohajerani, M. H., and McNaughton, B. L. (2018). Hippocampus-dependent emergence of spatial sequence coding in retrosplenial cortex. Proceedings of the National Academy of Sciences, 115(31):8015–8018.

Merre, P. L., Esmaeili, V., Charrière, E., Galan, K., Salin, P.-A., Petersen, C. C., and Crochet, S. (2018). Reward-Based Learning Drives Rapid Sensory Signals in Medial Prefrontal Cortex and Dorsal Hippocampus Necessary for Goal-Directed Behavior. Neuron, 97(1):83–91.e5.

Miller, A. M., Mau, W., and Smith, D. M. (2019). Retrosplenial Cortical Representations of Space and Future Goal Locations Develop with Learning. Current Biology, 29(12):2083–2090.e4.

Miller, A. M. P., Vedder, L. C., Law, L. M., and Smith, D. M. (2014). Cues, context, and long-term memory: the role of the retrosplenial cortex in spatial cognition. Frontiers in human neuroscience, 8:586.

Mitchell, A. S., Czajkowski, R., Zhang, N., Jeffery, K., and Nelson, A. J. D. (2018). Retrosplenial cortex and its role in spatial cognition. Brain and Neuroscience Advances, 2(3):239821281875709.

Mohan, H., Gallero-Salas, Y., Carta, S., Sacramento, J., Laurenczy, B., Sumanovski, L. T., Kock, C. P. J. d., Helmchen, F., and Sachidhanandam, S. (2018a). Sensory representation of an auditory cued tactile stimulus in the posterior parietal cortex of the mouse. Scientific Reports, 8(1):7739.

Mohan, H., Haan, R. d., Broersen, R., Pieneman, A. W., Helmchen, F., Staiger, J. F., Mansvelder, H. D., and Kock, C. P. J. d. (2019). Functional Architecture and Encoding of Tactile Sensorimotor Behavior in Rat Posterior Parietal Cortex. The Journal of Neuroscience, 39(37):7332–7343.

Mohan, H., Haan, R. d., Mansvelder, H. D., and Kock, C. P. d. (2018b). The posterior parietal cortex as integrative hub for whisker sensorimotor information. Neuroscience, 368:240–245.

Nikbakht, N. and Diamond, M. E. (2021). Conserved visual capacity of rats under red light. eLife, 10:e66429.

O’Connor, D. H., Clack, N. G., Huber, D., Komiyama, T., Myers, E. W., and Svoboda, K. (2010). Vibrissa-based object localization in head-fixed mice. The Journal of neuroscience: the official journal of the Society for Neuroscience, 30(5):1947–1967.

O’Keefe, J. (1976). Place units in the hippocampus of the freely moving rat. Experimental Neurology, 51(1):78–109.

O’Keefe, J. and Dostrovsky, J. (1971). The hippocampus as a spatial map. Preliminary evidence from unit activity in the freely-moving rat. Brain Research, 34(1):171–175.

Olcese, U., Iurilli, G., and Medini, P. (2013). Cellular and Synaptic Architecture of Multisensory Integration in the Mouse Neocortex. Neuron, 79(3):579–593.

Ottink, L., Hoogendonk, M., Doeller, C. F., Geest, T. M. V. d., and Wezel, R. J. A. V. (2021). Cognitive map formation through haptic and visual exploration of tactile city-like maps. Scientific Reports, 11(1):15254.

O’Connor, D. H., Krubitzer, L. E., and Sliman, J. B. (2021). Of mice and monkeys: Somatosensory processing in two prominent animal models. Progress in Neurobiology, 201:102008.

Pammer, L., O’Connor, D. H., Hires, S. A., Clack, N. G., Huber, D., Myers, E. W., and Svoboda, K. (2013). The Mechanical Variables Underlying Object Localization along the Axis of the Whisker. Journal of Neuroscience, 33(16):6726–6741.

Pereira, A., Ribeiro, S., Wiest, M., Moore, L. C., Pantoja, J., Lin, S.-C., and Nicolelis, M. A. L. (2007). Processing of tactile information by the hippocampus. Proceedings of the National Academy of Sciences, 104(46):18286–18291.

Petersen, C. C. H. (2007). The Functional Organization of the Barrel Cortex. Neuron, 56(2):339–355.

Pho, G. N., Goard, M. J., Woodson, J., Crawford, B., and Sur, M. (2019). Taskdependent representations of stimulus and choice in mouse parietal cortex. Nature communications, 9(1):2596.

Pluta, S. R., Lyall, E. H., Telian, G. I., Ryapolova-Webb, E., and Adesnik, H. (2017). Surround Integration Organizes a Spatial Map during Active Sensation. Neuron, pages 1–20.

Pnevmatikakis, E. a., Soudry, D., Gao, Y., Machado, T. A., Merel, J., Pfau, D., Reardon, T., Mu, Y., Lacefield, C., Yang, W., Ahrens, M., Bruno, R., Jessell, T. M., Peterka, D. S., Yuste, R., and Paninski, L. (2016). Simultaneous Denoising, Deconvolution, and Demixing of Calcium Imaging Data. Neuron, 89(2):285–299.

Poulter, S., Lee, S. A., Dachtler, J., Wills, T. J., and Lever, C. (2021). Vector Trace cells in the Subiculum of the Hippocampal formation. Nature neuroscience, 24(2):266–275.

Powell, A., Connelly, W. M., Vasalauskaite, A., Nelson, A. J. D., Vann, S. D., Aggleton, J. P., Sengpiel, F., and Ranson, A. (2020). Stable Encoding of Visual Cues in the Mouse Retrosplenial Cortex. Cerebral Cortex (New York, NY), 30(8):4424–4437.

Rice, F. L., Mance, A., and Munger, B. L. (1986). A comparative light microscopic analysis of the sensory innervation of the mystacial pad. I. Innervation of vibrissal follicle-sinus complexes. Journal of Comparative Neurology, 252(2):154–174.

Rivard, B., Li, Y., Lenck-Santini, P.-P., Poucet, B., and Muller, R. U. (2004). Representation of Objects in Space by Two Classes of Hippocampal Pyramidal Cells. The Journal of General Physiology, 124(1):9–25.

Save, E., Cressant, A., Thinus-Blanc, C., and Poucet, B. (1998). Spatial Firing of Hippocampal Place Cells in Blind Rats. The Journal of Neuroscience, 18(5):1818–1826.

Sikes, R. W., Vogt, B. A., and Swadlow, H. A. (1988). Neuronal responses in rabbit cingulate cortex linked to quick-phase eye movements during nystagmus. Journal of neurophysiology, 59(3):922–936.

Smith, J. B. and Alloway, K. D. (2010). Functional Specificity of Claustrum Connections in the Rat: Interhemispheric Communication between Specific Parts of Motor Cortex. Journal of Neuroscience, 30(50):16832–16844.

Sofroniew, N. J., Cohen, J. D., Lee, A. K., and Svoboda, K. (2014). Natural Whisker-Guided Behavior by Head-Fixed Mice in Tactile Virtual Reality. The Journal of Neuroscience, 34(29):9537–9550.

Stacho, M. and Manahan-Vaughan, D. (2022). Mechanistic flexibility of the retro-splenial cortex enables its contribution to spatial cognition. Trends in Neurosciences, 45(4):284–296.

Sugar, J., Witter, M. P., Strien, N. M. v., and Cappaert, N. L. M. (2011). The retros-plenial cortex: intrinsic connectivity and connections with the (para)hippocampal region in the rat. An interactive connectome. Frontiers in Neuroinformatics, 5:7.

Tsao, A., Moser, M.-B., and Moser, E. I. (2013). Traces of experience in the lateral entorhinal cortex. Current biology, 23(5):399–405.

Vann, S. D., Aggleton, J. P., and Maguire, E. A. (2009). What does the retrosplenial cortex do? Nature reviews. Neuroscience, 10(11):792–802.

Vedder, L. C., Miller, A. M. P., Harrison, M. B., and Smith, D. M. (2016). Retrosplenial Cortical Neurons Encode Navigational Cues, Trajectories and Reward Locations During Goal Directed Navigation. Cerebral Cortex, pages 1–11.

Weible, A. P., Rowland, D. C., Monaghan, C. K., Wolfgang, N. T., and Kentros, C. G. (2012). Neural Correlates of Long-Term Object Memory in the Mouse Anterior Cingulate Cortex. Journal of Neuroscience, 32(16):5598–5608.

Wijngaarden, J. B. v., Babl, S. S., and Ito, H. T. (2020). Entorhinal-retrosplenial circuits for allocentric-egocentric transformation of boundary coding. eLife, 9:e59816.

Witter, M. P., Doan, T. P., Jacobsen, B., Nilssen, E. S., and Ohara, S. (2017). Architecture of the Entorhinal Cortex A Review of Entorhinal Anatomy in Rodents with Some Comparative Notes. Frontiers in systems neuroscience, 11:46.

Wolbers, T., Klatzky, R., Loomis, J., Wutte, M., and Giudice, N. (2011). Modality-Independent Coding of Spatial Layout in the Human Brain. Current Biology, 21(11):984–989.

Wyss, J. M. and Groen, T. v. (1992). Connections between the retrosplenial cortex and the hippocampal formation in the rat: a review. Hippocampus, 2(1):1–11.

Zahler, S. H., Taylor, D. E., Wong, J. Y., Adams, J. M., and Feinberg, E. H. (2021). Superior colliculus drives stimulus-evoked directionally biased saccades and attempted head movements in head-fixed mice. eLife, 10:e73081.

Zhang, K., Ginzburg, I., McNaughton, B. L., and Sejnowski, T. J. (1998). Interpreting neuronal population activity by reconstruction: unified framework with application to hippocampal place cells. Journal of neurophysiology, 79(2):1017–1044.

Zhao, X., Wang, Y., Spruston, N., and Magee, J. C. (2020). Membrane potential dynamics underlying context-dependent sensory responses in the hippocampus. Nature Neuroscience, 23(7):881–891.

